# Decoding Huge Phage Diversity: A Taxonomic Classification of Lak Megaphages

**DOI:** 10.1101/2024.02.01.578382

**Authors:** Ryan Cook, Marco A. Crisci, Hannah V. Pye, Andrea Telatin, Evelien M. Adriaenssens, Joanne M. Santini

**Author notes:** Joint first authors (alphabetical order) (Ryan Cook, Marco A. Crisci, Hannah V. Pye).

## Abstract

High-throughput sequencing for uncultivated viruses has accelerated the understanding of global viral diversity and uncovered viral genomes substantially larger than any that have so far been cultured. Notably, the Lak phages are an enigmatic group of viruses that present some of the largest known phage genomes identified in human and animal microbiomes, and are dissimilar to any cultivated viruses. Despite the wealth of viral diversity that exists within sequencing datasets, uncultivated viruses have rarely been used for taxonomic classification. We investigated the evolutionary relationships of 23 Lak phages and propose a taxonomy for their classification. Predicted protein analysis revealed the Lak phages formed a deeply branching monophyletic clade within the class *Caudoviricetes* which contained no other phage genomes. One of the interesting features of this clade is that all current members are characterised by an alternative genetic code. We propose the Lak phages belong to a new order, the “Grandevirales”. Protein and nucleotide-based analyses support the creation of two families, three sub-families, and four genera within the order “Grandevirales”. We anticipate that the proposed taxonomy of Lak megaphages will simplify the future classification of related viral genomes as they are uncovered. Continued efforts to classify divergent viruses are crucial to aid common analyses of viral genomes and metagenomes.

## Introduction

Advancements in metagenomic sequencing have uncovered phage genomes greater than 200 kb (designated jumbo phages), and megaphages with genomes between ∼500 kb and ∼735 kb [1]. The colloquially named Lak phages are a group of large dsDNA phages which were first identified in human gut metagenomes from Laksam Upazila, Bangladesh [2]. Genomes related to these megaphages have since been detected in international gut metagenomes from humans and various animals including pigs, non-human primates, horses and tortoises [2, 3]. To date, 23 phylogenetically related Lak-like genomes (∼476-660 kb) have been resolved to completion [2, 3]. However, the Lak phages remain uncultured. CRISPR-spacer targeting has indicated that Lak phages infect bacteria in the genus *Prevotella*, which are highly abundant in the gut microbiome of humans and animals that consume high-fibre and low-fat diets [2, 4]. The absence of integrases in the Lak phage genomes and no evidence of prophages in metagenomes suggests that these are virulent phages, rather than temperate phages that integrate into the bacterial host genome [3].

Although Lak phages are associated with gut microbiomes, some of the largest known complete phage genomes (∼630-735 kb) were assembled from aquatic environment samples [1, 5]. For example, Mar_Mega_1 (650 kb) was recently assembled and found to be widespread across global marine samples [5]. Phylogenetic analyses of the Lak-like megaphages indicates that they form a single clade when compared to other dsDNA phages, with the marine megaphages belonging to a sister clade of the Lak phages. Mar_Mega_1 was proposed to represent a novel family, and forms a clade with LR756502 (642 kb) and LR745206 (635 kb), which were identified in freshwater metagenome samples from France and Japan, respectively [5].

Lak megaphages have previously been predicted to use an alternative genetic code, which differs to that commonly used by bacterial species and most other known phages (genetic code 11), whereby the TAG stop codon is repurposed to encode glutamine (genetic code 15) [6]. However, phage recoding cannot be validated without evidence of functional protein translation [3, 7, 8]. Moreover, the phenomenon of stop-codon repurposing has been reported for ∼5% of phage genomes recovered from human and animal gut metagenomes, and is suggested to prevent premature production of late-stage proteins and possibly regulate lysis [6]. Notably, some members of the *Crassvirales* are also thought to repurpose their stop codons [6, 9, 10]. However, this feature is not a characteristic of all *Crassvirales,* and only seems to occur within specific families such as the *Suoliviridae* [10]. Whether alternative code use is indicative of all megaphages remains unknown.

Beyond their shared re-purposing of TAG codons, there are several parallels between the Lak megaphages and *Crassvirales.* First identified from human metagenomic data in 2014, the original crAssphage was thought to be a ubiquitous resident of the human gut [11]. Subsequently, an increasing number of crAss-like phages were identified from metagenomic data and shown to be highly abundant in the GI tract of adults that consume low-fibre and high-fat diets; promoting *Bacteroides*-rich enterotypes instead of *Prevotella*-rich enterotypes [12, 13]. Despite their widespread abundance, isolation of the first crAss-like phage from *Bacteroides intestinalis* (ΦcrAss001) occurred in 2018 [14]. Since then, the order *Crassvirales* has been ratified by the ICTV, an additional member has been cultured (ΦcrAss002), and the particle structure of ΦcrAss001 has recently been resolved [15, 16]. *Crassvirales* therefore offer an example of how analysis of uncultured viral genomes can lead to novel biological insights of ecologically significant viruses.

While the classification of prokaryotic viruses traditionally followed conservation of their morphology, this has been superseded by genomic-based approaches [17]. The introduction of a 15-rank virus hierarchy (species to realm) has most notably led to the abolition of the order Caudovirales (tailed phages), with all members reassigned to the class *Caudoviricetes* [17, 18]. This restructuring now enables megaphages to be assigned to higher taxonomic ranks, following ICTV guidelines alongside the four principles of establishing a universal viral taxonomy [19]. Therefore, the aim of this study was to investigate the evolutionary history of Lak phages and define taxa for the 23 Lak megaphages using genomic-based methods.

## Methods

### Genomes

Complete phage genomes that had been previously resolved from metagenomic analysis of the gut microbiomes of humans [2, 3], baboons [2, 20], horses [3], dogs [21, 22] and pigs [3] were downloaded from ggKbase (University of California, Berkley) (Supplementary Table 1), and additional large unclassified phage genomes were extracted from the INPHARED database (February, 2023; Supplementary Table 2) [23]. Emboss v6.6.0 was used to determine the length of each phage genome and the GC content (Supplementary Table 1 & 2) [24].

### Predicted Proteome Analysis

A set of publicly available phage genomes (n=3,539) belonging to the realm *Duplodnaviria*, which comprises all tailed bacteriophages, was extracted from the INPHARED database (February, 2023) [23], and the up-to-date taxonomy for each genome was extracted from the Virus Metadata Resource (VMR; https://ictv.global/vmr). The classified members of the *Duplodnaviria* were combined with the genomes used in this study for input with the standalone version of ViPTree v1.1.2 [25], and the output was visualised with IToL [26]. To determine the potential effect of alternative TAG codon usage on coding capacity, open reading frames (ORFs) were predicted on all genomes using Prodigal v2.6.3 with both translation table 11 and 15 [27]. Coding capacity was calculated as the sum length of predicted ORFs as a percentage of total genome length.

### Core Genome Analysis

Open reading frames (ORFs) were predicted using Prodigal-gv v2.11.0-gv, a modified version of Prodigal that predicts alternative codon usage and has been optimised for gene identification on virus genomes (https://github.com/apcamargo/prodigal-gv) [27–29]. The translated ORFs were clustered using MMseqs2 v13.45111 at 70% identity with the --cluster-mode 1 flag [30]. Sequences from each cluster were compared to publicly available MMseqs2 profiles of the PHROGs database (https://phrogs.lmge.uca.fr/downloads_from_website/phrogs_mmseqs_db.tar.gz) using mmseqs search with -e 1E-05 and the best hit per ORF was retained [30, 31]. The most frequent PHROG hit was used to infer the function of the cluster. Presence/absence of proteins clusters were plotted using seaborn v0.12.2 [32].

Alignments were produced for the 72 protein clusters found on the 22 genomes belonging to the proposed family “Lakviridae” using MAFFT v7.520 [33], and the alignments were used as input for IQ-TREE v2.2.2.3 with -B 1000 and -m TEST [34]. As the IQ-TREE “-m TEST” flag optimises models for individual alignments before combining into a final model, multiple models were used in the final tree (Supplementary Table 3). The resulting concatenated protein maximum likelihood phylogenetic tree was visualised using IToL [26].

### Intergenomic Similarity of Lak Phages

The intergenomic similarity of the 23 Lak phage genomes was deduced by multiplying the average nucleotide identity (ANI) with the aligned fraction of the genome for each pair of genomes. ANI was determined using fastANI v1.33 [35] for all genomes using the many-to-many parameter, for multiple reference and query genomes. The aligned fraction was calculated by dividing the number of fragments that were aligned as orthologous matches between each genome by the total number of sequence fragments for each genome, both of which were computed within the fastANI output. The ANI value was then multiplied by the aligned fraction to compute the intergenomic similarity. The average intergenomic similarity between genomic pairs was visualised in R v4.2.2 using pheatmap v1.0.12 [36].

## Results

This study included 56 phage genomes > 200 Kb that are not currently classified to any taxonomic rank, including 23 so-called Lak phages (Supplementary Table 1). For the purposes of this study, only complete genomes were included. The 56 genomes ranged from 203-735 kb in length and 24-55 molGC (%), with the 23 Lak phages ranging from 476-660 kb and 26-31 molGC (%).

The 56 genomes included in this analysis all belonged to prokaryotic viruses with dsDNA genomes which were predicted to encode the set of proteins that are characteristic of the newly ratified class *Caudoviricetes.* These features include putative tail proteins, a major capsid protein with the HK97 fold, a portal protein, and the terminase large subunit. Based on the presence of these proteins, all 56 phages are automatically assigned to the class *Caudoviricetes* and this analysis therefore sought to further classify these phages from the rank of order to species. To infer the order and family rank, a proteomic approach was used. The 56 genomes were processed using VipTree alongside all currently classified members of *Caudoviricetes* (n=3,539; February 2023), producing a hierarchically clustered tree based on pairwise tBLASTx scores (Figure 1). The 23 Lak phages formed a deeply branching monophyletic clade that contained no other genomes, with Mar_mega_1 being the only genome in its nearest sister clade (Figure 1A). Of the remaining 32 large phage genomes, 14 were interspersed with known genomes belonging to sister clades of the Lak phages, and the remaining 18 were very distant on the tree (Figure 1A).

**Figure 1.**
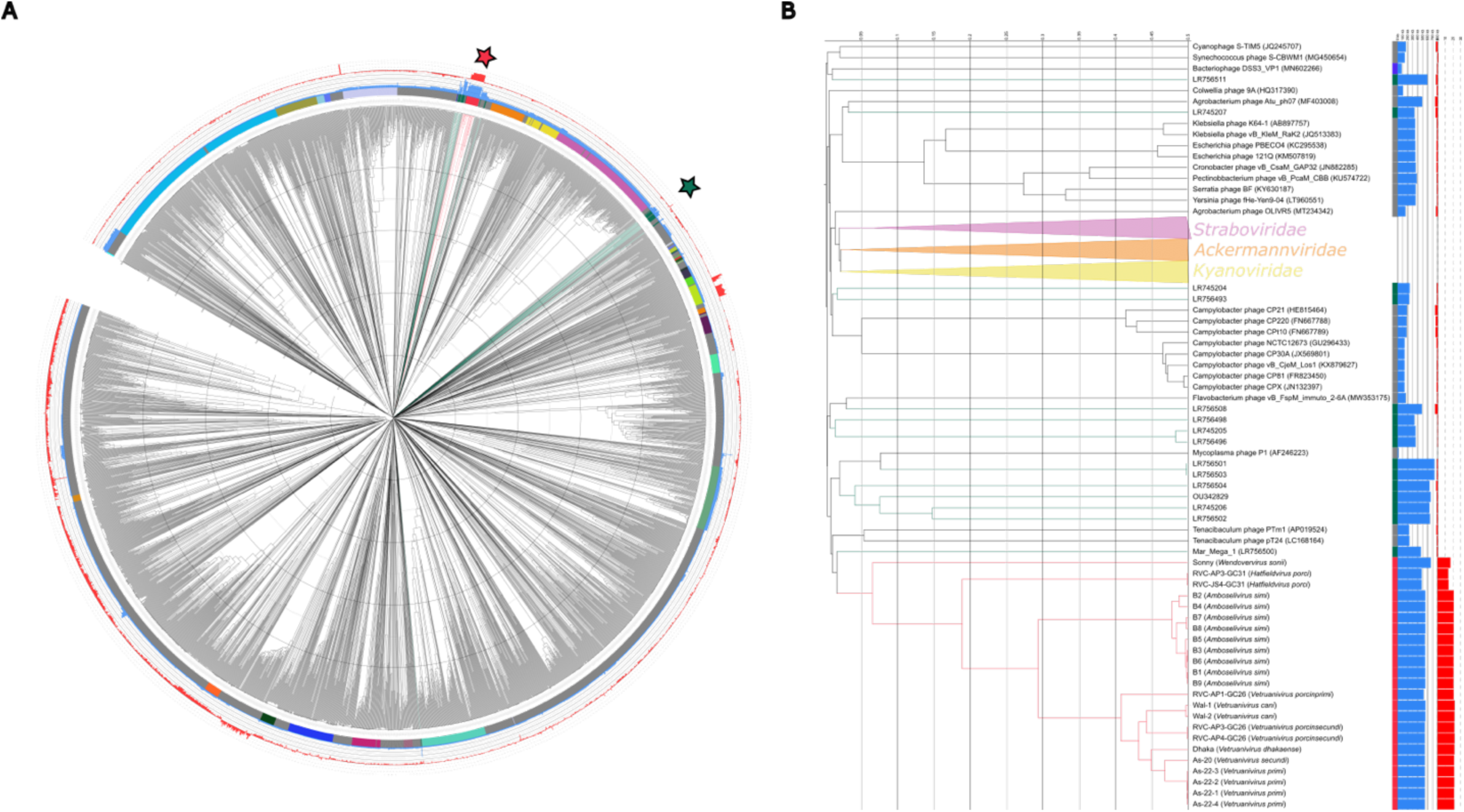
Proteomic tree of Megaphages amongst currently classified *Duplodnaviria*. (**A**) ViPTree proteomic tree of “megaphages” and *Duplodnaviria* with viral family shown in the coloured ring. Blue bar chart (inner) represents genome length and red bar chart (outer) shows difference in coding capacity when using translation table 15 rather than 11 (i.e., coding capacity of 90% using 15 and 70% using 11 would lead to a difference of 20). The “Grandevirales” members are shown with red branches and other megaphages are shown with green and highlighted by a star with corresponding colour. (**B**) A pruned tree showing the “Grandevirales” with nearest sister clades only. The distances shown in ViPTree were calculated from genomic distances based upon normalised tBLASTx scores and the tree was rooted at the mid-point.

Lak phages are thought to re-purpose TAG codons to encode glutamine, rather than a stop codon, as this is observed elsewhere in biology [37]. To determine if this feature is unique to and conserved within Lak phages, we predicted ORFs on the 56 genomes used in this analysis and the 3,539 representative members of the realm *Duplodnaviria* using both translation table 11 (standard bacterial) and table 15 (re-purposed TAG). The mean coding capacity of the 23 Lak phages with translation table 11 was 69% (SD ± 1.9), and increased to 89% (SD ± 1.2) when using translation table 15 (Figure 1B; Supplementary Table 4). This feature was conserved among all Lak phages but was not observed in Mar_Mega_1 (table 11 94%, table 15 93%). Considering the distance to other phages, and conserved alternative codon use amongst this clade, we propose the Lak phages represent a new order of phages and suggest the name “Grandevirales”. As the 23 proposed members of “Grandevirales” were highly divergent from other phages used in this study, only these 23 were carried forward for taxonomic classification.

To infer taxonomy at and below the rank of family, we performed a core protein analysis on the proposed members of “Grandevirales”. The 23 members of “Grandevirales” shared no core proteins across all genomes at 70% amino acid sequence identity (Figure 2). The genome of phage Sonny (OR769223) was a clear outlier with the predicted proteome vastly different from the remaining 22 phages. Only when the threshold for amino acid identity was lowered to 25% did this phage genome share one core gene with the 22 others. We therefore suggest that this phage represents a new family and propose the name “Epsomviridae”. The other 22 “Grandevirales” shared 72 core proteins at 70% amino acid identity, the vast majority of which have unknown functions (Figure 2; Supplementary Table 5 & 6). Of 1,623 unique protein clusters within this group, there was a median of 583 proteins per genome and the 72 core proteins represent a mean average of 12.5% (SD ± 0.8%). We propose that these phages belong to a second new family, “Lakviridae”. The proposed “Lakviridae” consist of two clear groupings that are distinct from one another at a higher level than genus (Figure 2). Therefore, we suggest the “Lakviridae” is divided into two subfamilies, “Quadringentisvirinae” and “Quingentivirinae”. Encouragingly, this pattern is congruent with that of the VipTree analysis performed for higher level classification (Figure 1).

**Figure 2.**
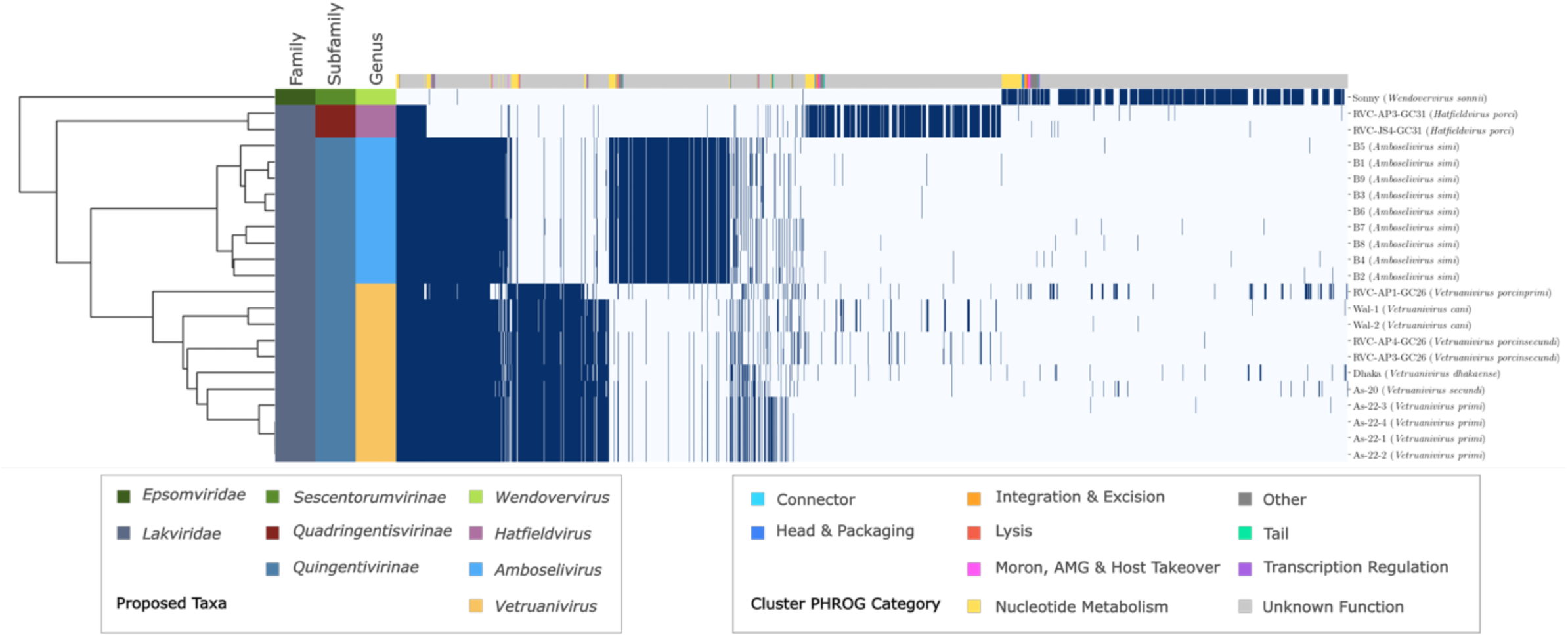
Shared protein clusters for phages of “Grandevirales”. Heatmap showing presence/absence of proteins clustered at 70% identity with the dendrogram showing hierarchical clustering. Colour strips on the y-axis show proposed taxonomy, and colour strips on the x-axis show predicted function of the protein cluster derived from PHROGs. X-axis labels show strain name with proposed species name in brackets.

The 72 proteins core to the proposed “Lakviridae” were aligned and used as input for a concatenated protein phylogeny. This analysis revealed two deeply branching clades that mirror the results of the core genome analysis, lending further support to the creation of subfamilies “Quadringentisvirinae” and “Quingentivirinae” (Figure 3). The proposed “Quingentivirinae” form two distinct clades that we suggest represent two separate genera (Figure 3). One clade, the “Amboselivirus”, consists of the nine highly similar “Lak B” phages that were identified from baboon samples (Figure 3). The other clade, “Vetruanivirus”, consists of 11 genomes identified from separate studies of human and pig samples (Figure 3).

**Figure 3.**
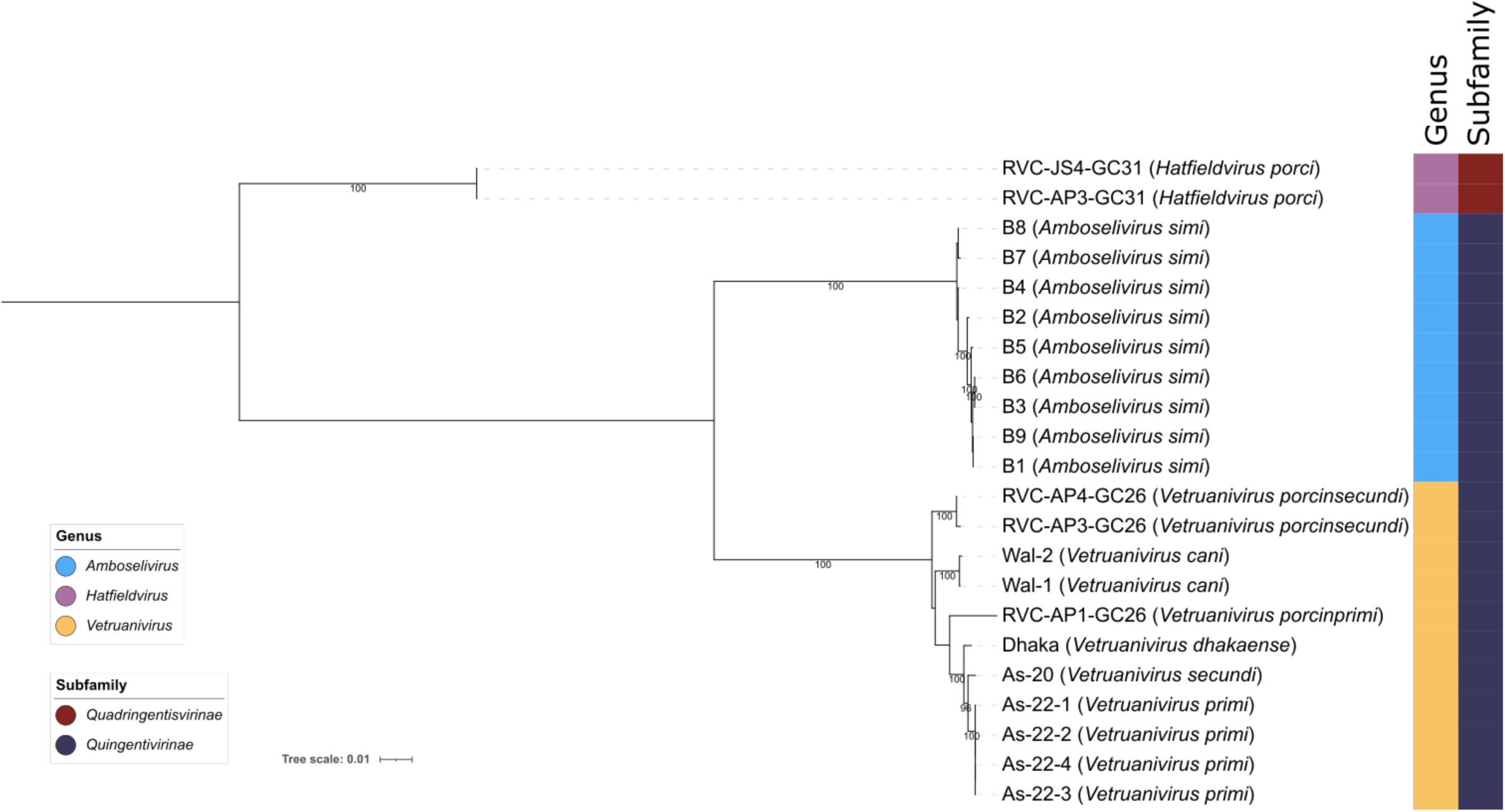
Core genome phylogeny of “Lakviridae” phages. A concatenated protein phylogeny of translated sequences of 72 “core” genes present on the 22 members of the proposed family “Lakviridae”. Proposed species names for each strain are indicated in brackets. Alignments were performed using MAFFT, and the tree was produced using IQ-Tree with 1000 rapid bootstraps and -m TEST to optimize model fits for each alignment. Tree is rooted at the midpoint and bootstraps ≥ 95% are shown. The coloured strips indicate proposed genera and subfamilies. Node labels are based on strain names with proposed species names shown in brackets.

To elucidate species level phylogenetic relationships, we investigated the intergenomic similarity of the proposed “Grandevirales”. The 23 “Grandevirales” phages formed four distinct clusters with varying intergenomic similarity (Figure 4). Three of the genomes (As-22-2, As-22-1 and As-22-4) shared almost 100% identity (Figure 4; Supplementary Table 7). Four phages were classified into the species “Vetruanivirus primi”, due to an intergenomic similarity score of >95%. According to the same criteria, two phages were assigned to each species of “Vetruanivirus porcinsecundi” and “Vetruanivirus cani”. Eleven phages were classified as the genus “Vetruanivirus” as they were all within the typical genus demarcation of 70%, as defined by the ICTV. Another set of nine phages, all isolated from the Baboon gut (B1-B9), made up the sole species (“Amboselivirus simi”) of the currently proposed “Amboselivirus” genus (Figure 4; Supplementary Table 7). Almost all of these Baboon-associated phage genomes had a pairwise identity of >95%, and the three genomes with marginally less than 95% pairwise identity (B4 compared to B1, and B4 compared to B9) still cluster distinctly with other members of the “Amboselivirus” genus within 95% identity (Figure 4), hence why we chose to classify them into the same species.

**Figure 4.**
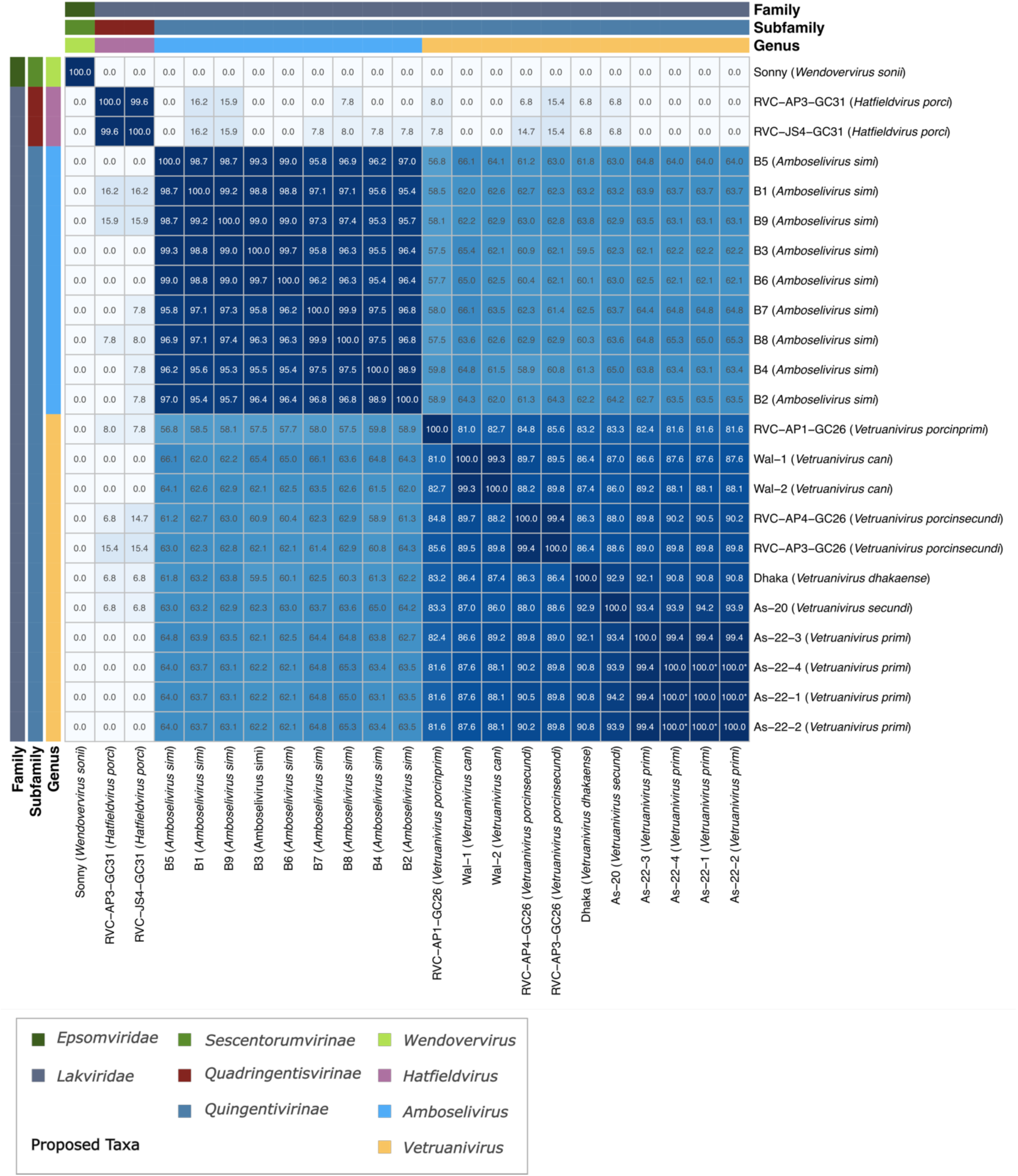
Intergenomic similarity of “Grandevirales” phages. Heatmap showing intergenomic distance (ANI multiplied by aligned fraction) of “Grandevirales” phages with proposed taxonomy shown in coloured strips. Those shown as having a similarity of 100.0* are highly similar but not identical with a low number of single nucleotide variations between them. Axis labels are based on strain names with proposed species names shown in brackets.

Only two phages formed the “Hatfieldvirus porci” species (∼99.36% intergenomic similarity) within the “Hatfieldvirus” genus. Interestingly, members of this genus share little similarity (< 20%) with members of the “Amboselivirus” and “Vetruanivirus” genera (Figure 4; Supplementary Table 7). The sole member of the “Wendovervirus sonii” species shares no detectable similarity with any other phages included in the analysis, and hence further supports our reasoning to designate this phage as a member of a separate family (“Epsomviridae”) (Figure 4; Supplementary Table 7). All strain names used in the current study have remained consistent with the nomenclature used in the respective publications [2, 3].

The computational analysis of the previously published 23 Lak-like megaphage genomes has led to the proposed formation of the order “Grandevirales”, encompassing two families, “Lakviridae” and “Epsomviridae”, three sub-families and four separate genera (Table 1).

**Table 1.**
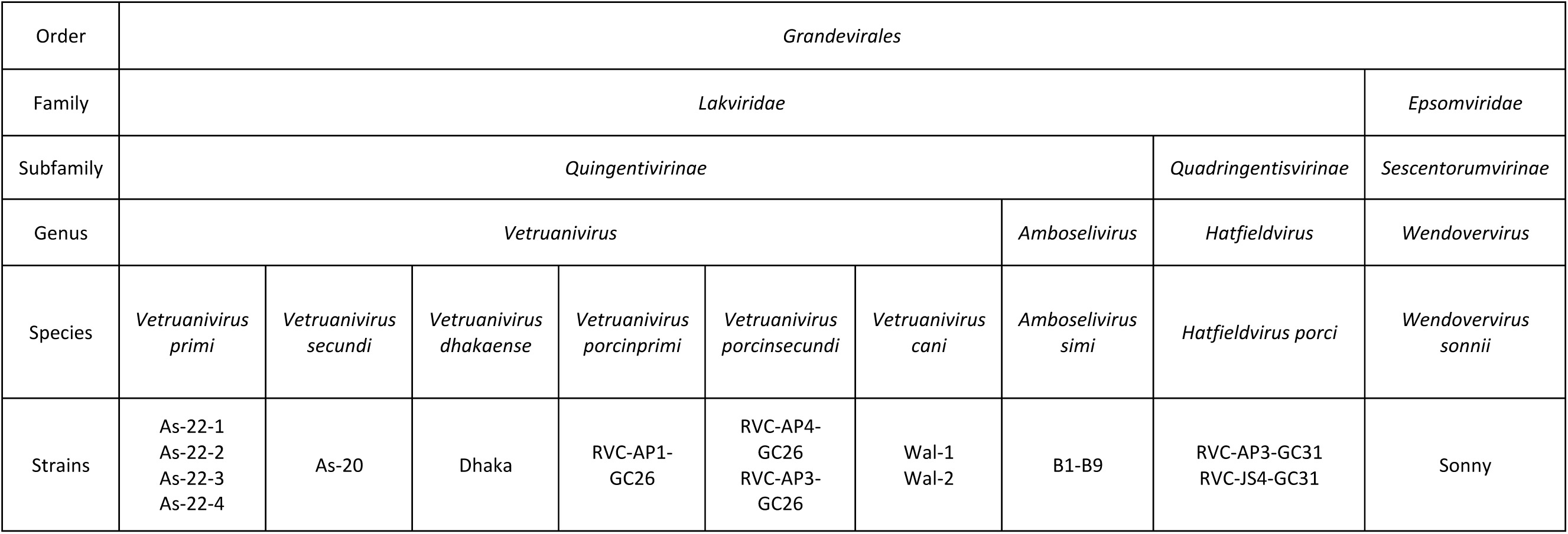
Proposed taxonomic classification of Lak-like megaphages.

### Rationale and Justification of Taxonomic Names

All taxonomic names proposed in this manuscript follow the guidance suggested by Postler *et al*., (2022), with all proposed species using a latinised binomial name [38].

#### Order “Grandevirales”

The name “Grandevirales” is proposed for the order, as Grande means large (or great) in Latin and in many other European languages. Based on the present analyses, some related phages in this order have genomes under 500 Kb, therefore it would be unsuitable to denote the order as ‘Mega’, given that ‘megaphage’ has historically been used to describe phages with genomes over 500 Kb [2]. Furthermore, the order *Megavirales* has been used to describe large eukaryotic viruses (>100 Kb) [39]. Given that all phages within this order are ‘huge’, i.e., > 200 Kb, and that the order encompasses the largest known complete phage genomes, the order name “Grandevirales” was chosen.

#### Family “Lakviridae”

The family “Lakvirdae” was selected as the founding members of this family (A1 and A2) were all identified from the gut microbiomes of people consuming arsenic-contaminated water in Laksam Upazila, Bangladesh [2].

#### Family “Epsomviridae”

The second family proposed in this study is “Epsomviridae”, named after Epsom, Surrey (UK). Epsom is a town famous for horse racing and is the location where the racehorse “Sonny” was stabled and trained. In the current proposal, only one megaphage belongs to this family, which was identified from the horse gut microbiome [3]. Sonny was therefore used for the species and strain name.

#### Subfamilies “Quingenti-”, “Quadringenti-” and “Sescentorum-” -virinae

Subfamily names were chosen based on genome sizes of the founding members of each subfamily. ‘Quingenti’, ‘Quadringentis’, and ‘Sescentorum’ are Latin for 400, 500 and 600, respectively. For “Quingentivirinae”, founding members have genome lengths within the 500 Kb range. For “Quadringentisvirinae”, current members have ∼476 Kb genomes, and one phage has currently been classified into the subfamily “Sescentorumvirinae”, with a ∼660 Kb genome. Although the genome lengths of founding members have been used as the basis of subfamily names, this is not a criterion for taxonomic classification and members of each subfamily form distinct clusters within “Lakviridae” and “Epsomviridae”.

#### Genus “Vetruanivirus”

The genus name “Vetruanivirus” is an amalgamation of the words Veterinary and Eruani, which encompasses the isolation source of current phages in this genus. Some phages were isolated from pigs at the Royal Veterinary College (RVC), hence “*Vet-*“, whilst other strains were isolated from individuals living in a village called Eruani in Laksam, Bangladesh, hence “ruani-“ which forms “Vetruanivrus”.

#### Genus “Amboselivirus”

Phage genomes belonging to the proposed “Amboselivirus” genus (B1-B9) were resolved from faecal samples collected from Kenyan yellow baboons living in the Amboseli national park. The current sole species has been given the name “Amboselivirus simi” (‘simia’ is Latin for primate/monkey, with simi in the genitive form). Strains within the proposed genus “Amboselivirus” (B1-B9) are maintained as described in the original publication [2].

#### Genus “Hatfieldvirus”

Two of the phage genomes identified from pig samples formed the “Hatfieldvirus” genus, which has been named according to the sampling location. These phages were identified from faecal samples from pigs reared and cared for at the RVC in Hatfield, Hertfordshire (UK). The sole species name has been given as, “Hatfieldvirus porci”, as porci is Latin for pig. Strains of this proposed genus (and those predicted to be placed in it), have so far only been found in pig gastrointestinal tracts.

#### Genus “Wendovervirus”

The sole genome belonging to this genus, Sonny, was assembled from sequencing data of a microbiome sample from a horse stabled at the Wendover stables in Epsom [3].

## Discussion

An increase in the number of high-quality curated phage genomes has transformed phage taxonomy, and morphology-based classification has been superseded with robust genomic frameworks [17, 40, 41]. In this study, 23 Lak phages were taxonomically classified via comprehensive pangenome analysis, concatenated protein phylogeny, and analysis of their intergenomic similarity. The taxonomy and analysis of these Lak phages – which we can now call grandeviruses – posed several challenges related to their origin, alternative codon usage and large genome sizes.

The megaphage genomes classified in this study were resolved from metagenomes and the phages themselves remain uncultured, due to the difficulty in isolating these large phages from biological samples, as also described with *Crassvirales* [15, 16]. Phage cultivability is dependent on identification of the bacterial host and optimal growth conditions, neither of which can be easily determined from metagenomes despite a plethora of tools developed to aid in the identification of these phage-host pairings, such as iPHoP (which uses RaFAH, WIsH, oligonucleotide frequencies, PHP, and BLAST), HostPhinder and PHERI [42–50]. Alternative methods to isolate phages with large genomes have emerged, and suggest the use of filters with pore sizes >0.2 µm, and decreasing the concentration of the overlay agar used [51].

Pangenomes are often used to classify bacterial taxa according to the presence of core and accessory genes in bacterial strains. During the current study, multiple pangenome construction tools, including roary [52], Panaroo [53] and ggcaller [54], were used to generate a pangenome of the 23 megaphages and related huge phages. However, we found that all of these tools were inappropriate for this particular group of sequences. Many published tools do not support translation table 15, the stop codon reassignment that is suggested in these phages. Furthermore, we found no available tools that could construct the pangenome of sequences that use different translation tables to one another (i.e., determining the pangenome of megaphages that use translation table 15, alongside their nearest relatives that use translation table 11). Therefore, a manual method for elucidating megaphage pangenomes was devised that used prodigal-gv to account for the alternative codon usage [28].

While the pangenome tools developed for bacteria were not compatible with the alternative codon use of the “Grandevirales” phages, the virus tools struggled with their genome sizes. The sheer size of the megaphage genomes in question (476-660 Kb) made it difficult to use phage and virus-specific published tools to determine viral intergenomic similarity, such as VIRIDIC [55] and VirClust [56]. We therefore manually implemented the general methodology used by VIRIDIC to compute intergenomic similarity, ensuring that the nucleotide identity was normalised according to the aligned fraction of the genome to avoid exaggerated intergenomic similarity scores [55]. Interestingly, there was a low percentage (6.8-16.2%) of intergenomic similarity observed between phages belonging to different genera. This warrants further investigation using comparative genomics to elucidate the origin of these stretches of sequence identity, whether they are the result of recombination or horizontal gene transfer between phages or the bacterial host.

We were able to make a few observations with potential biological relevance that could be topics of further investigation. All grandeviruses were identified from gut microbiomes of humans and mammals associated with humans (evolutionarily or through close contact). Phages belonging to each species were isolated from the same source, for example all nine strains of “Amboselivirus simi” were discovered in baboon gut microbiomes, reflected in their chosen species name. In contrast, at the genus rank multiple microbiome hosts were observed for the genus “Vetruanivirus”. Further identification and classification of these phages in microbiomes and hopefully isolation in culture will answer questions about their function in the gut and their niche-adaptation in different animals.

## Conclusion

The results presented here, combined with previously published work, provide robust evidence that the Lak phage clade is monophyletic compared to other known phage genomes and justifies the creation of a new viral order, “Grandevirales”. The order encompasses some of the largest phage genomes ever reported and can be further sub-divided into two new families and three subfamilies according to concatenated protein phylogeny and intergenomic similarity. Four novel genera have also been proposed, encompassing 23 phage strains. Overall, this study has overcome the challenges associated with the classification of “megaphages”, has successfully classified phages resolved from metagenomes, and provided justification for the classification of huge viral genomes based on a shared core genome, intergenomic similarity and phylogeny.

## Acknowledgments

Not applicable.

## Authors’ Contributions

Conceptualisation: MAC, EMA, JMS; Data curation: RC, MAC, HVP; Formal analysis: RC, MAC, HVP, AT; Funding acquisition: JMS; Supervision: EMA, JMS; Writing – Original Draft Preparation: RC, MAC, HVP, AT, EMA, JMS; Writing – Review & Editing: EMA, JMS.

## Availability of Data and Materials

The genome sequences analysed in this study are from previous studies and are publicly available. Details and accession numbers can be found in Supplementary Tables 1 & 2.

## Ethical Approval and Consent to Participate

Not applicable.

## Consent for Publication

Not applicable.

